# DeepGANnel: Synthesis of fully annotated single molecule patch-clamp data using generative adversarial networks

**DOI:** 10.1101/2020.06.25.171918

**Authors:** Sam T. M. Ball, Numan Celik, Elaheh Sayari, Lina Abdul Kadir, Fiona O’Brien, Richard Barrett-Jolley

**Affiliations:** Faculty of Health and Life Science, University of Liverpool, Liverpool, United Kingdom

## Abstract

Development of automated analysis tools for “single ion channel” recording is hampered by the lack of available training data. For machine learning based tools, very large training sets are necessary with sample-by-sample point labelled data (e.g., 1 sample point every 100microsecond). In an experimental context, such data are labelled with human supervision, and whilst this is feasible for simple experimental analysis, it is infeasible to generate the enormous datasets that would be necessary for a big data approach using hand crafting. In this work we aimed to develop methods to generate simulated ion channel data that is free from assumptions and prior knowledge of noise and underlying hidden Markov models.

We successfully leverage generative adversarial networks (GANs) to build an end-to-end pipeline for generating an unlimited amount of labelled training data from a small annotated ion channel “seed” record, and this needs no prior knowledge of theoretical dynamical ion channel properties. Our method utilises 2D CNNs to maintain the synchronised temporal relationship between the raw and idealised record. We demonstrate the applicability of the method with 5 different data sources and show authenticity with t-SNE and UMAP projection comparisons between real and synthetic data. The model would be easily extendable to other time series data requiring parallel labelling, such as labelled ECG signals or raw nanopore sequencing data.

## 1. Introduction

Ion channels are transmembrane proteins that control the flow of ions from one side of the membrane to the other; they play a fundamental role in the function of key biological processes, such as the generation and propagation of action potentials (Hodgkin & Huxley, 1952; Neher & Sakmann, 1976). Due to their fundamental role in biological processes, dysfunction of ion channels can lead to a number of conditions (known as “channelopathies”), including cystic fibrosis (Quinton, 1990; Welsh, 1990). Beyond being directly responsible for health conditions, ion channels are also targeted for other uses, such as pesticides in agriculture (Silver et al., 2014) or therapeutic drugs for other conditions such as pain management (Skerratt & West, 2015) and a number of neuropsychiatric disorders (Imbrici, Conte Camerino & Tricarico, 2013).

The broad roles of ion channels motivates research into their function. The development of the Nobel Prize winning patch-clamp electrophysiological technique (Hamill, Marty, Neher, Sakmann & Sigworth, 1981; Sakmann & Neher, 1984) allowed for “real time” recording of the function of different ion channels to changes of conditions, such as the application of drugs. In particular, single channel recording has the ability to capture a single ion channel opening and closing in real-time, and provides researchers with the tools necessary to investigate the mechanisms behind the function of these proteins.

Analysis of patch clamp single channel data is notoriously difficult, typically taking skilled researchers significant time to manually supervise. Many different models have been developed to attempt to solve this problem (Celik et al., 2020; Gnanasambandam, Nielsen, Nicolai, Sachs, Hofgaard & Dreyer, 2017a; Hotz et al., 2013; Qin, 2004) – however most of these methods are developed using simulated data generated by simulating a continuous time hidden Markov model (HMM), then applying a layer of filtered Gaussian noise on top (for example, Anderson, Ermentrout & Thomas, 2015; Colquhoun, Hawkes & Srodzinski, 1996; Gillespie, 1977; Nicolai & Sachs, 2013; Voldsgaard Clausen, 2020). Whilst this method can generate a large amount of data for training, the data itself may have hidden biases (including a user encoded HMM gating schema), since it is entirely dependent on a priori assumptions about underlying model and characteristic noise dynamics.

Generative Adversarial Networks (GANs) (Goodfellow et al., 2014) have proven to be important for generative tasks, such as image generation (Brock, Donahue & Simonyan, 2019; Karras, Laine & Aila, 2019) or audio synthesis (Donahue, McAuley & Puckette, 2019). GANs have been used in electrophysiology to create new unsynchronised datasets (Qin et al., 2020; Truong, Kuhlmann, Bonyadi, Querlioz, Zhou & Kavehei, 2019), with the former example being used to generate data for model training, but continuous labelling of data (millisecond-by-millisecond) has remained an unsolved challenge. GANs have great potential to allow a data driven approach to synthesis, using a small amount of “real”, lab recorded data to generate an unlimited amount of simulated data; the advantage of this method is that noise and artefacts present in the real training dataset also appear in the generated data.

In this work we develop a GAN model to generate realistic, simulated ion channel data from existing patch-clamp recordings from several channel phenotypes, each with a continuous parallel state label. These outputs could then be used to build more accurate ion channel analysis models. We use a further series of analysis tools to evaluate the model’s performance – including T-SNE and UMAP projections to investigate if each channel’s high dimensional characteristics are reflected in the simulated data.

## 2. Methods

### 2.1 Model design

We propose a GAN based model to generate synthetic time-series data that includes realistically similar features to real ion channel molecule currents. Figure 1 describes the complete pipeline of the proposed DeepGANnel model in this work. The architecture of the GAN model introduced in this work is following the regular DC-GAN, but applies the convolutional neural networks that demonstrated efficiently to produce time series datasets in previous works (Delaney, Brophy & Ward, 2019; Zhu, Ye, Fu, Liu & Shen, 2019). In the following sections, the utilized neural networks briefly are covered along with the processed data information and evaluation metrics criteria.

**Fig. 1.**
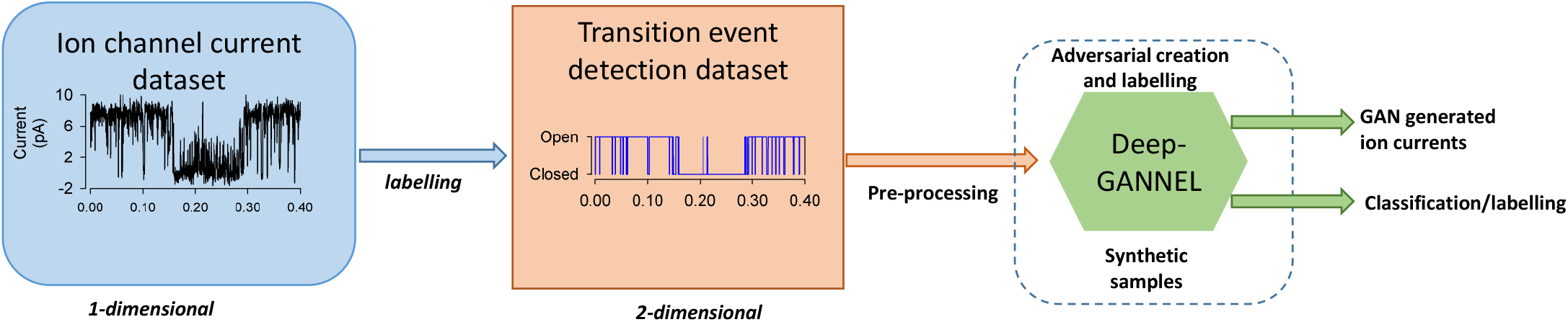
Pipelines for DeepGANNEL Development. Data are recorded using standard electrophysiological techniques. These raw data are then labelled using existing software and expert supervision. Pre-processing makes copies of these data adding a small amount of gaussian noise to the raw data and reshapes the two signals into 2D “images” (2×1280).

### 2.2 Generative Adversarial Networks (GAN)

A GAN typically consists of two neural networks competing against each other; the generator network tries to convert random noise into observations that seem as if they have been sampled from the original data; while the discriminator network aims to classify whether the sampled data comes from the original dataset or output of the generator network by predicting a class probability. The training process is employed in an adversarial manner between the two networks by updating the parameters of both models based on updates of the discriminator; the generator is looking to maximise the discriminator’s uncertainty in classifying an output as real or fake, and the discriminator is looking to minimise this. A more detailed, mathematical formulation of these loss functions, along with the formulation of the constituent layers of these models can be found in Appendix 1, and the full model architecture of both generator and discriminator including shapes are included in Appendix 2 and 3 respectively.

### 2.3 Design of the Networks

The DeepGANnel networks are based on the successful DC-GAN model (Radford, Metz & Chintala, 2015), with the input and output shapes of the generator changed to images with dimensions *n* by *m* (one channel for the raw signal, one for the idealisation, *n* and *m* were typically 2 and 1280, but this numbers can easily be varied depending on context) along with changes to the subsequent hyperparameters along with pre/post processing to facilitate this change.

The architecture of the generator *G* consists of strided deconvolutional layers that allow the network to undergo spatial learning with its own up-sampling; batch normalisation layers allow for stabilising learning parameters by normalising inputs and Leaky ReLU activation functions for all layers, except for the *Tanh* function that is used in the output layer. The input to this model is a matrix of latent noise that is transformed into the output signal, and the output of the model is a two-dimensional sample generated from the noise that can be sent to the input of the discriminator for training the model.

The discriminator *D*, is also a deep convolutional neural network. Similar to the generator, this model combines strided convolution layers to downsample the input data to obtain a binary classification of the input record (real or generated). In our discriminator architecture, the batch normalisation layers are not used, but instead a dropout regularised technique was used at each layer. The last convolution layer in our *D* network is flattened and passed into a sigmoid function for classification, but otherwise similarly to the generator function;Leaky ReLU activation functions are used for all other layers. Figure 2 shows the architecture of our DeepGANnel model in this work including both *G* and *D* networks. The Adam optimiser was used initially with 0.0001 learning rate for initial training process. Manual tuning of the learning rate parameters was needed during training to avoid overfitting from one of the models.

**Fig. 2.**
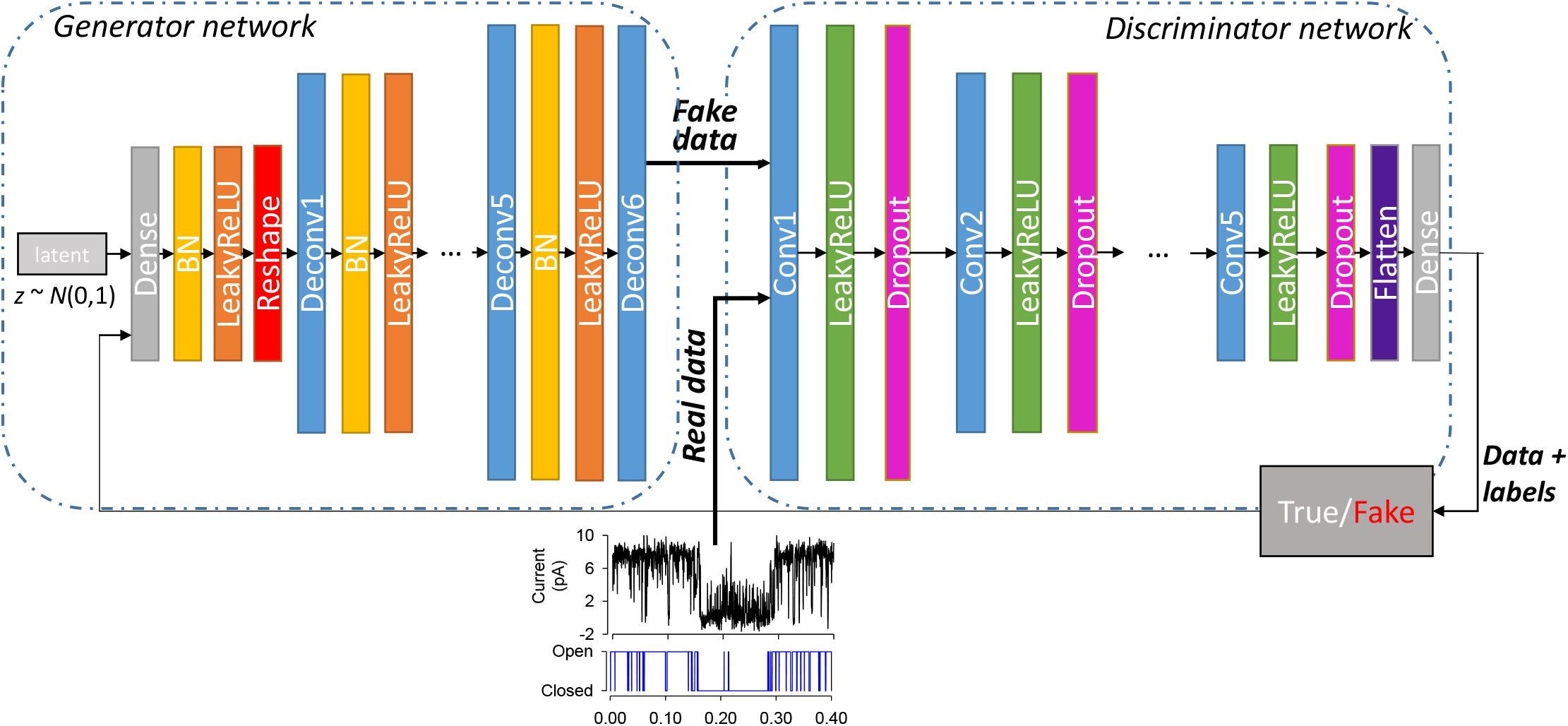
DeepGANnel network architecture. On the left is a generator network of DeepGANNEL in which the input is a random noise and output is a 2D image by 2×1280 that is converted from 1-D time series. The generator model in this architecture uses three deconvolution layers (upsampling) to produce an image from seed (random noise shape by 2×1280). Several Dense layers are used to take the noise input and transform into a desired image by up-sampling (Deconv or transposed convolutional layer) steps size of 2×1280. At each layer, Leaky Rectified Linear Unit (LeakyReLU) is used as activation function with batch normalisation, except output layer which uses tanh activation function. The discriminator model comprises of several convolutional layers with the same activation function (LeakyReLU) at each layer to classify whether an image is real or fake by comparing the sampled data to real data. Dropout layers were also appended to all convolutional layers except input layer with the value of 0.3 to reduce overfitting. Finally, a Dense output layer is used after flattening the network structure with classified labels.

### 2.4 Data sources

Ion channel recordings were obtained from cell-attached patch clamp electrophysiology on a number of different cell types; for detailed methods please see the associated references.

In the case of tracheal chondrocytes (“Phenotype C”), methods similar to (Lewis *et al*. 2013), except that trachea was isolated, cut into small pieces, and thence treated as for articular chondrocytes. In all cases, animals were previously euthanised for unassociated reasons; no animals were killed or harmed for this study. Typically, patch pipettes were fabricated using 1.5Cmm *o*.*d*. borosilicate glass capillary tubes (Sutter Instrument, USA, supplied by INTRACEL, UK). They were pulled using a two-step electrode puller (Narishige, Tokyo, Japan) and when filled with recording solutions, had a resistance of approximately 5-10CMΩ. In each case, data was recorded using cell-attached patch clamp with an Axopatch 200/a amplifier (Axon Instruments, USA). Low-pass filtering was set to 1⍰Hz and data were digitized at 5 kHz with a Digidata 1200A interface or CED 1401 (CED, Cambridge, UK). Recordings were made with WinEDR (John Dempster, University of Strathclyde, UK).

In addition, in order to achieve a diverse population of ion channel records to test our GAN model, we simulated ion channel data using the simulation feature in WinEDR (John Dempster, University of Strathclyde, UK) using default rate constants (“Phenotype B”). Idealisation/annotation of raw data was performed with QuB (Nicolai & Sachs, 2013).

### 2.5 Evaluation metrics

GANs are considered successful when they implicitly learn the distribution of samples of the real dataset. We assess the efficiency of the proposed DeepGANnel model to simulate single molecule data by comparing real to GAN simulated data using a number of different approaches. The standard metrics for GANs are the so-called generator loss and discriminator loss. These are calculated as the logistic binary cross entropy loss – this is calculated as:

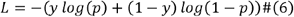

Where *y* is the true label and *p* the predicted label. For the generator this is calculated once, with the true label being whether the output is fake or not, and the predicted label being whether the discriminator predicted if the output is fake. For the discriminator, this is calculated twice then totalled – once for the fake output as above, and again in the same manner for real outputs.

We also use other methods to measure model success. Although there is currently some debate as to which metrics are most appropriate for measuring GAN performance, we will utilize a two-sample test called maximum mean discrepancy (Gretton, Borgwardt, Rasch, Schölkopf & Smola, 2008), and a classical evaluation metric for time series defined as dynamic time warping (Sakoe & Chiba, 1978) to evaluate model performance.

We also use the t-SNE and UMAP dimensionality reduction algorithms as a means to compare and contrast the different ion channels and sources in a more meaningful way.

#### Maximum Mean Discrepancy

The maximum mean discrepancy (MMD) measures the dissimilarity between two probability distributions *p*_*r*_ and *p*_*g*_ – one from the real data distribution and one from the GAN respectively, by comparing statistics of the samples. In general, given a kernel *K* : *X* x *Y* → *R*, and samples 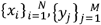 an estimation of MMD is equated as follows:

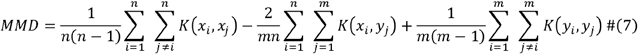

The smaller MMD measurement, the greater similarity between the distributions. A modified Python library’s (Tensorflow) two-sample test (Gretton, Borgwardt, Rasch, Scholkopf & Smola, 2012) was used to determine MMD using the Gaussian kernel for the above calculation.

#### Dynamic Time Warping

Dynamic time warping (DTW) is a traditional calculation for measuring the dissimilarity between two groups of time series data. DTW warps the series temporally to minimise the Euclidean distance between points and then calculates the distance between the two warped records. The warping of one signal occurs using the following method. An N by M matrix D is constructed where N is the length of the first signal x and M the length of the second signal y. Starting at *D*_0,0_, the matrix is filled out iteratively using the following formula:

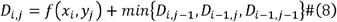

While *I* = 1,…,*N* and *j =* 1,…., *M*. This matrix can then be used to find the most appropriate warping for the data, for which then a Euclidian similarity metric can be used. Due to the large collection and individual size of simulated ion channel records that was used in this work, a dedicated library (FastDTW (Salvadora & Chan, 2007)) was utilized to approximate DTW.

#### T-SNE and UMAP

Dimensionality reduction algorithms allow us to visualise high dimensional data in a low dimensional format, typically within a visualisation such as a scatter plot. Principle component analysis (PCA), for example, reduces the dimension of data by choosing a new orthogonal basis for the data based on the maximal variance of the data. For non-linear datasets, this dimensionality reduction may not accurately convey the shape or pattern of the higher dimensional data, as points that are close together in Euclidian space but far away in the context of the data are brought together during the PCA transformation. The so-called “kernel trick” can solve this problem by providing a non-linear transformation to the space before the PCA algorithm is applied, however this requires that this non-linear transformation to be found algorithmically, which can be time consuming and inaccurate. The t-SNE (Van Der Maaten & Hinton, 2008) algorithm focuses on local similarity rather than global similarity by considering only a neighbourhood around each point in high dimensional space. The algorithm first constructs a matrix of probability distributions for each pair of points in the higher dimensional space such that close points have a high probability and far away points have a low probability. The algorithm then constructs a second matrix of probability distributions for the lower dimensional space and minimises the Kullback-Leibler divergence between the two via gradient descent. This method preserves local similarities, but global similarities are lost between far away points. Uniform Manifold Approximation and Projection (UMAP) (McInnes, Healy & Melville, 2018) was developed to attempt to maintain the global similarities by using the same general method as t-distributed stochastic neighbour embedding (T-SNE) (the construction of two matrices and optimisation to fit the lower dimensional one to the higher dimensional one), but uses a topological transformation of the data and a different divergence metric to achieve stronger global similarity. Note that in both cases we trained these projections on the super dataset together (i.e., all raw and GAN records concatenated together). We used the standard Python packages t-SNE from Scikit Learn and UMAP from the umap-learn conda-forge. To give an objective indication of whether there was a significant difference between manifold clusters between original and GAN simulated data, we tested for statistical difference between each pair with the R package ClusterSignificance (Serviss, Gådin, Eriksson, Folkersen & Grandér, 2017).

### 2.6 Computing platform

A Nvidia Titan X GPU with 12 GB of RAM was used for the experiments to train and generate realistic ion channel molecule current records alongside the event classifications of generated records using the DeepGANnel model. The GAN models were built in Tensorflow 2.x (including Keras) using Jupyter notebooks as a user-friendly interface exploiting the GPU. With this computing platform and software, trained models Once trained, the included code on this equipment generates 0.4 seconds of data (ie. one 4096×2-datapoint “record” in under 4ms).

## 3. Results

Single (ion channel) molecule datasets were recorded using the patch-clamp technique and our standard protocols, and approximately 30 seconds was recorded under constant conditions (room temperature etc.). The resulting datasets were annotated (idealised) in QuB to produce a two-dimensional signal (dimension 1 = raw signal, dimension 2 = continuous annotation). Following robust scaling (Scikit learn) and reshaping these data were passed to the DeepGANnel model for approximately 10,000 epochs. Processing within each training epoch included augmentation with an invisibly small amount of Gaussian noise (approximately 0.1% of signal amplitude) applied to each data window. Figure 3A shows the characteristic evolution of discriminator and generator losses. Typical generator losses were 10x or more the discriminator loss. The first 250 epochs of the more sophisticated MMD and DTW losses are shown in Figure 3B. Figure 3 C and D show examples of raw (real) input data and a representative strip of post train DeepGANnel generated data. The two first analyses that are typically conducted in patch-clamp research are amplitude histograms and kinetic analysis. In addition to the objective metrics, amplitude histograms are shown in Figures 3E (real events) and Figure 3F (GAN simulated events), are also clearly similar in terms of size and shape.

**Fig. 3.**
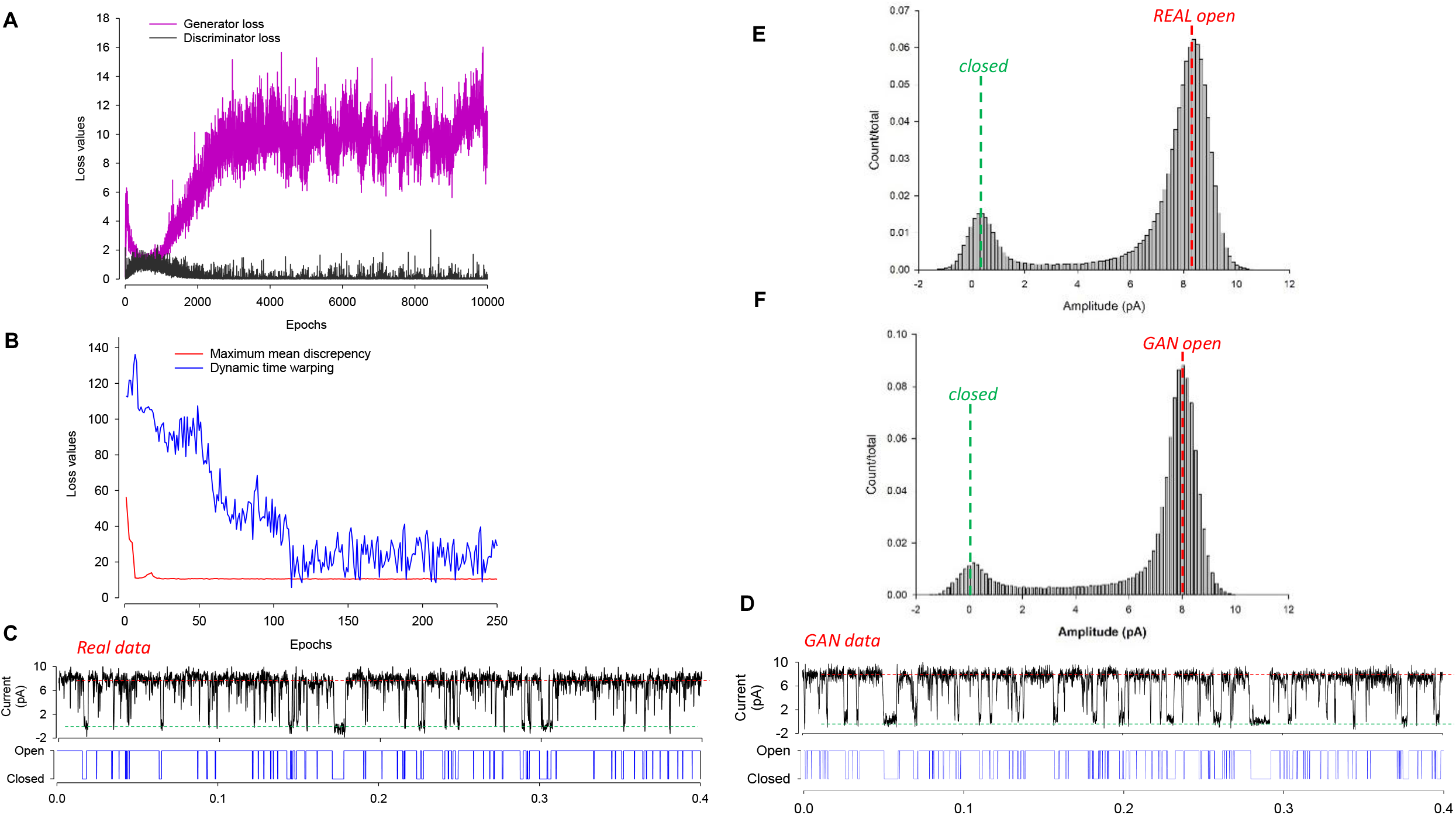
Realistic labelled single molecule data are synthesised by DeepGANnel. (A) Discriminator and Generator losses plotted per epoch during training. (B) The maximum mean discrepancy (MMD) and dynamic time warping (DTW) evolution over the first 250 epochs as they approach equilibrium. (C and D) Examples of real (C) and GAN synthesised (D) labelled data records. (E and F) All points amplitude histograms calculated from the real (E) and GAN synthesised (F) data.

For an in-depth analysis we conducted full kinetic analysis of both raw (real molecular data) and GAN simulated data. These are shown in Figure 4, and there are similarities and differences between the real and GAN events. In terms of closed times, it is apparent that whilst the over-all distribution is similar between real (Figure 4A) and GAN (Figure 4C) there are differences. The real data included some long closed-events that are absent from the GAN simulated equivalent. In terms of open times again the over-all distribution is similar between real (Figure 4B) and GAN simulated (Figure 4D), but there is attenuation of the very long open sojourns in the GAN simulated data.

**Fig. 4.**
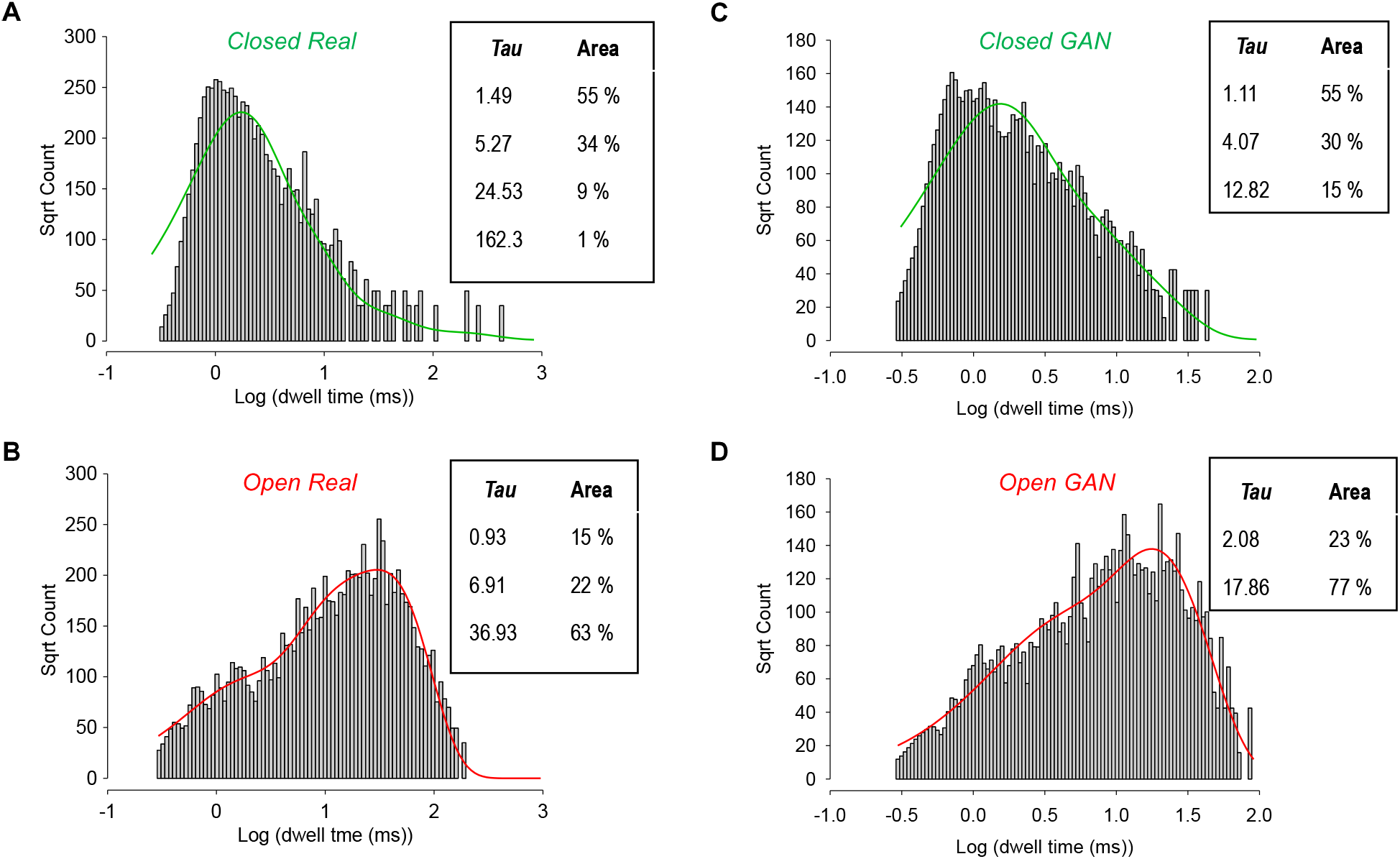
Comparative kinetic analyses of real and DeepGANnel simulated data. (A and B) Kinetic analysis from the real (input) single molecule data with closed times (A) fit with 4 exponentials and open times (B) fit with 3 exponentials. The respective time constants and weights are displayed in the inset tables. (C and D) Kinetic analysis from the GAN simulated (output) single molecule data with closed times (C) fit with 3 exponentials and open times (D) fit with 2 exponentials. The respective time constants and weights are again displayed in the inset tables.

To further test the ability of the GAN method to simulate a wide range of ion channel-like data we then trained DeepGANel on 4 further ion channel phenotypes (see Table I). Each of these is shown in Figure 5, where raw data, including the timeseries label, is shown a above the GAN simulated data. It is clear that each phenotype of channel is well represented by the matching GAN. To investigate more objectively how similar each GAN generated dataset is from its parent raw data we performed t-SNE and UMAP dimension reduction comparisons between each of the 5-phenotypes of data (Figure 6). It is clear that there is a tendency for each GAN manifold cluster (t-SNE or UMAP) to align closely with its parent real data projection. Following this, we systematically analysed the cluster separation between real and GAN clusters, but first, we investigated whether this method could distinguish between each real vs real combination, and each GAN vs GAN combination. We focussed on UMAP projections. Table II shows the comparisons between UMAPs of the real data. Each real dataset is significantly different to each other, demonstrating the power of UMAP cluster analysis to objectively classify real world data. Secondly, the equivalent analysis focusses on UMAP comparisons of the 5 different GAN generated datasets is show in Table III. In this case, all GAN simulated datasets are different from each other, except for the A and C datasets (different types of cartilage ion channels), which are not significantly different from each other. Finally, Table IV shows that cluster-analysis of UMAP projections is unable to see a statistically significant difference between the canine articular ion channel data (Phenotype A) and its respective GAN, the tracheal ion channel data (Phenotype C) and its respective GAN or the equine cartilage data (Phenotype E) and its derived GAN data. However, the GANs produced from channel phenotypes B and D were significantly distinguishable from their parent datasets (Phenotypes B and D).

**Table I.** Sources of raw data for development and testing of the GAN network.

| Code | Channel source | Methods Reference | Figure |
| --- | --- | --- | --- |
| Phenotype A | Canine articular chondrocyte | (Lewis et al., 2013) | 5A, 6A |
| Phenotype B | WinEDR simulated | N/A | 5B, 6B |
| Phenotype C | Tracheal chondrocyte | None, see below | 5C, 6C |
| Phenotype D | Paraventricular nucleus of the Hypothalamus | Feetham, Nunn, Lewis, Dart, & Barrett-Jolley, 2015) | 5D, 6D |
| Phenotype E | Equine articular chondrocytes | (Mobasheri, Lewis, Maxwell, Hill, Womack & Barrett-Jolley, 2010), | 1-4 |

**Table II.** Statistical separation of UMAP clusters between real datasets. Maximum *p-values* calculated by the ClusterSignificance permutation package in R (Serviss, Gådin, Eriksson, Folkersen & Grandér, 2017). *p-values* are given after at least 1000 permutations. Phenotype codes described in the methods, Table I.

|  | Phenotype A | Phenotype B | Phenotype C | Phenotype D | Phenotype E |
| --- | --- | --- | --- | --- | --- |
| Phenotype A | 1 | 0.02 | 0.001 | 0.001 | 0.001 |
| Phenotype B | - | 1 | 0.02 | 0.001 | 0.001 |
| Phenotype C | - | - | 1 | 0.001 | 0.001 |
| Phenotype D | - | - | - | 1 | 0.001 |
| Phenotype E | - | - | - | - | 1 |

**Table III.**
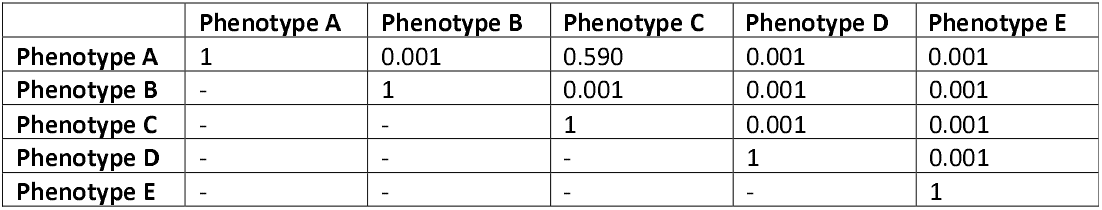
Statistical separation of UMAP clusters between GAN generated datasets. Maximum *p-values* calculated by the ClusterSignificance permutation package in R (Serviss, Gådin, Eriksson, Folkersen & Grandér, 2017). *p-values* are given after at least 1000 permutations. Phenotype codes described in the methods, Table I.

|  | Phenotype A | Phenotype B | Phenotype C | Phenotype D | Phenotype E |
| --- | --- | --- | --- | --- | --- |
| Phenotype A | 1 | 0.001 | 0.590 | 0.001 | 0.001 |
| Phenotype B | - | 1 | 0.001 | 0.001 | 0.001 |
| Phenotype C | - | - | 1 | 0.001 | 0.001 |
| Phenotype D | - | - | - | 1 | 0.001 |
| Phenotype E | - | - | - | - | 1 |

**Table IV.** Statistical separation of UMAP clusters between each real dataset and its GAN simulated equivalent. Maximum *p-values* calculated by the ClusterSignificance permutation package in R (Serviss, Gådin, Eriksson, Folkersen & Grandér, 2017). *p-values* are given after at least 1000 permutations.

| Phenotype | Source | p-val |
| --- | --- | --- |
| A | Canine articular chondrocyte | 0.739 |
| B | WinEDR simulated | 0.001 |
| C | Tracheal chondrocyte | 0.371 |
| D | Paraventricular nucleus of the Hypothalamus | 0.001 |
| E | Equine articular chondrocytes | 0.993 |

**Fig. 5.**
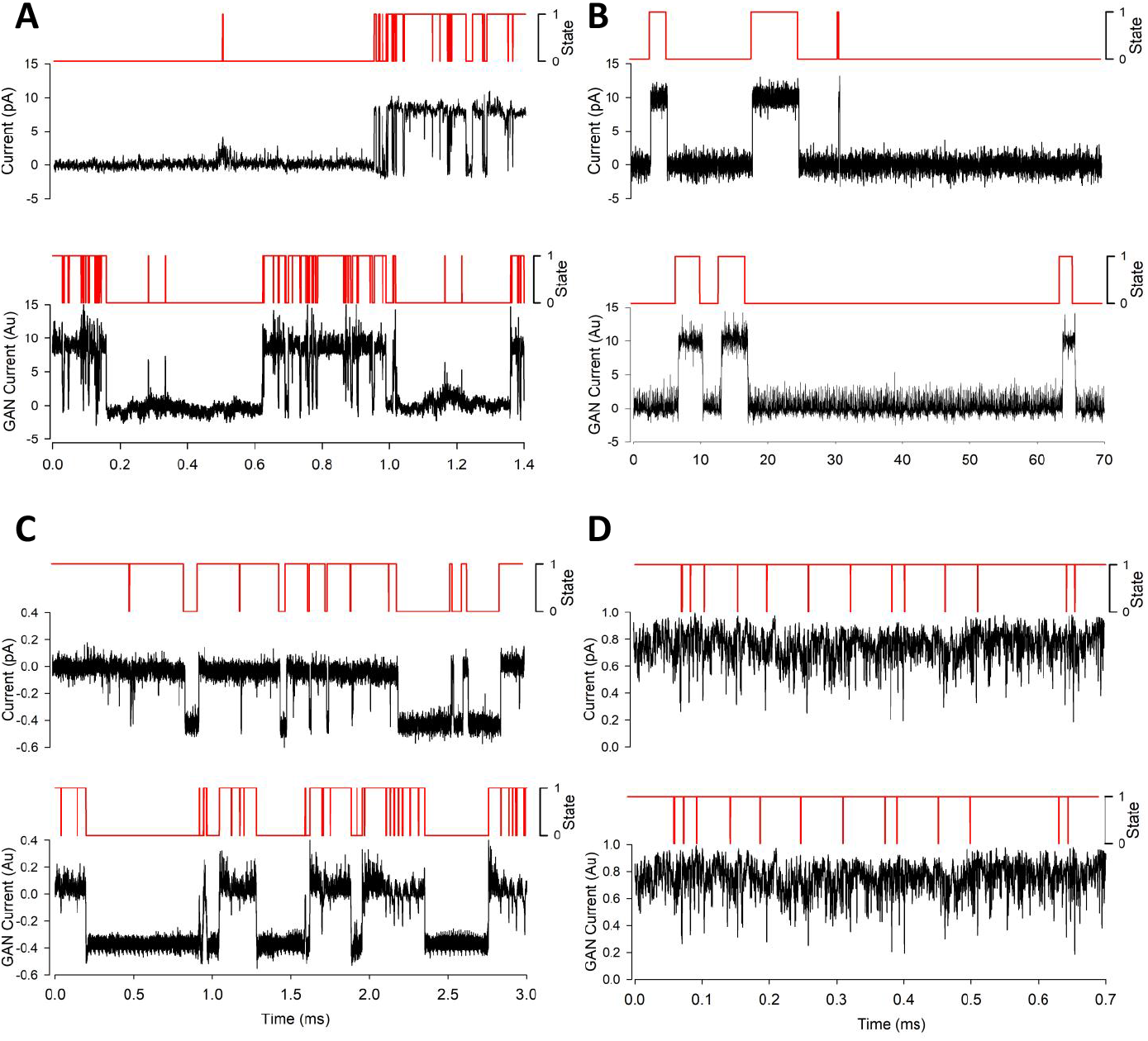
Sample data from both real dataset and DeepGANnel model output from numerous ion channels. In each subpanel, real raw and labelled data is displayed on top with simulated data below. (A): “Channel phenotype A” canine articular chondrocyte sample data and model output. (B): “Channel phenotype B”, WinEDR simulated sample data and model output. (C): “Channel phenotype C” tracheal chondrocyte sample data and model output. (D): “Channel phenotype D” PVN sample data and model output.

**Fig. 6.**
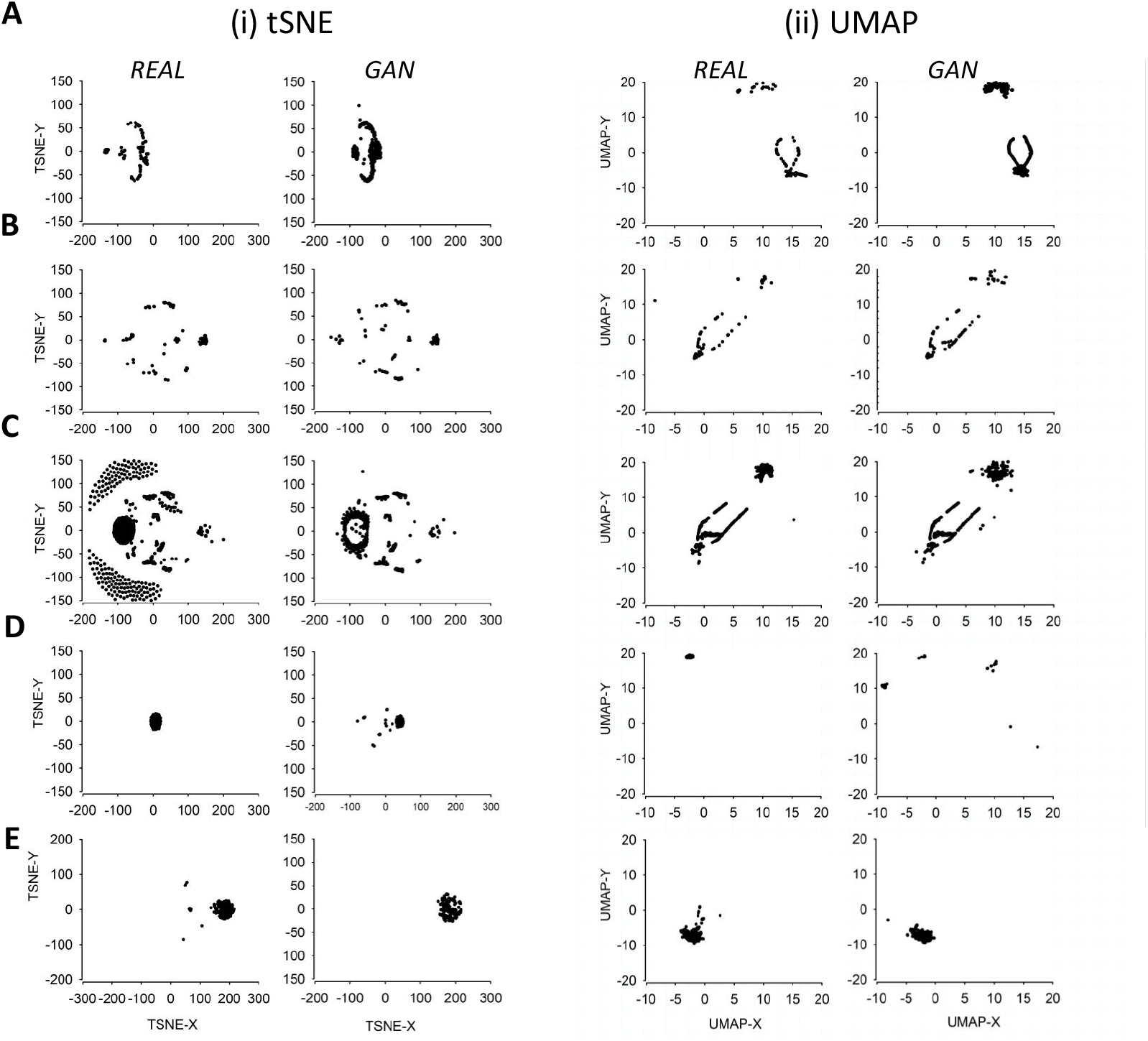
T-SNE and UMAP dimensional reduced visualisations for DeepGANnel and real data. For each of the channel phenotypes shown earlier (A to E), (i) T-SNE and (ii) UMAP projections are shown with “GAN” data on the left and “real” source data on the right. Sets of projections A to D correspond to those channel phenotypes shown in Figure 5, (A): “Channel phenotype A” canine articular chondrocyte. (B): “Channel phenotype C” WinEDR simulated. (C): “Channel phenotype B”, tracheal chondrocyte. (D): “Channel phenotype D” PVN data and (E) shows the same arrangement with the equine chondrocyte data shown in Figures 1 to 4.

## 4. Discussion

In this work, we have generated synthetic raw single-molecule timeseries data along with continuous synchronised annotation/idealisation using a generative adversarial network (GAN) based on both real ion channel single molecule data from cultured chondrocytes. We demonstrate that the GAN generated raw ion channel data was similar to those obtained by real ion channel data. We assessed success of the GAN by three methods and in each they proved successful, but retain some limitations.

A central problem in single molecule, including “ion channel” research is that analyses of data is laborious and frequently requires a degree of expert hand crafting to complete. The first step in such analysis is idealisation of the record, or in machine learning terms, annotating or labelling. Each time point (of which there will be many million) needs to be annotated as to how many molecule pores are open at that instant. This then becomes, effectively, a two-dimensional representation of the data. Currently, we are working with simple datasets with one type of molecule in the dataset, but in the future such analysis will extend this to more complex datasets. Clearly new analysis methods will also be necessary to get the maximum amount of information from complex single molecule data. A number of tools have been developed to address these issues (Colquhoun, Hatton & Hawkes, 2003; Gnanasambandam, Nielsen, Nicolai, Sachs, Hofgaard & Dreyer, 2017b; Juette et al., 2016; Nicolai & Sachs, 2013), including our own Deep-Channel deep learning model (Celik et al., 2020). For further development of similar or enhanced tools there is a lack of available training data. There are two clear choices for such data; (i) real biological data that has been annotated in some way or (ii) synthetic datasets. Both approaches have biases and severe limitations. Real data cannot be perfectly labelled; the real ground truth is unknowable and so will only be an approximation. In practical terms, the longer the length of the real data, the greater the “ground truth” errors will be. As a result, new machine learning methods will learn the errors of the existing technology. Furthermore, only simple datasets can be annotated and so this sets an upper limit on the complexity of the datasets that could be analysed by potential new tools. However, synthetic datasets also contain many biases and limitations, some of which may be entirely unanticipated or recognised. Therefore, the starting problem that our work addressed here was to create very large datasets of ion channel single molecule activity that could be used to develop single molecule analysis tools. We chose to investigate if GAN technology could provide a useful alternative source of data. In principle this would have the advantages of synthetic data in that one could produce unlimited amounts, but still retain nuance and subtle authenticity missed by mere simulation. We are also hopeful that such synthetic data could be used in development of ion channel modelling software and may allow for a novel type of data inference, extracting critical features in datasets that may be overlooked by traditional analysis. Visual inspection, and analysis of our similarity metrics demonstrates that DeepGANel can reconstruct faithfully simulate a number of different ion channel phenotypes. Clearly the datasets are not identical (that would be simply a copy), but raises the question of whether the simulation is *good enough* to be useful. To examine this, we show in Figure 7 an example experiment. We train our Deep-Channel model to analyse/idealise the data from phenotype C (Table IV, Figure 5C). Half of the small length of original raw data (75k datapoints equivalent to approx. 8 seconds of recording) is clearly insufficient to allow Deep-Channel to learn the appropriate features and idealise the remaining half of this dataset (Figure 7A, B, C). However, DeepGANel, once trained can produce any amount of data similar to this original dataset (See Figure 5C). We then trained Deep-Channel on this GAN dataset, 5x the size of the original, and performance is now acceptable (Figure 7D, E, F).

**Fig. 7.**
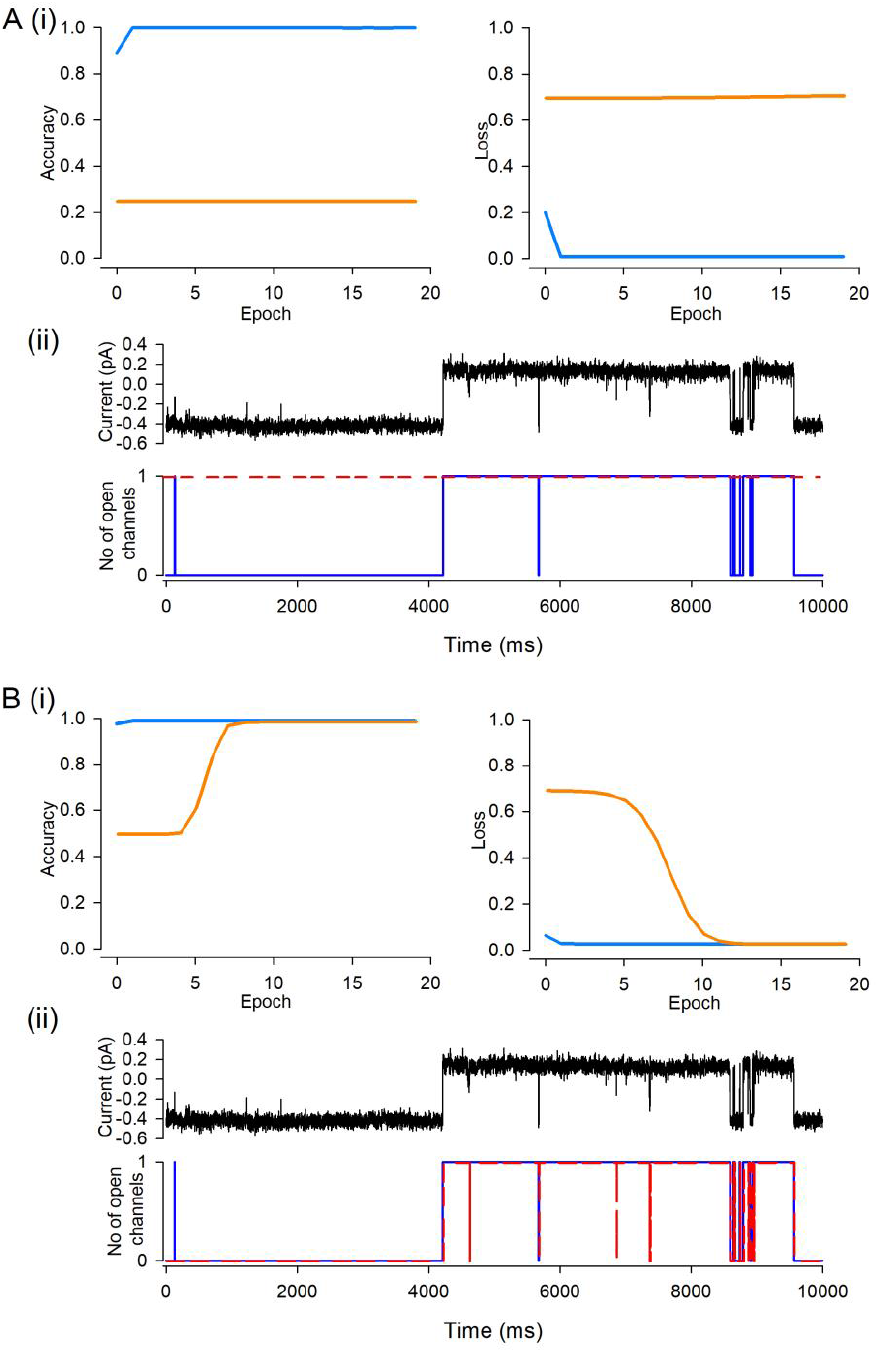
Potential of DeepGANnel to facilitate Deep Learning Training on Physiological time series data. This demonstration uses data from ion channel Phenotype C. The original raw data is 150k sample points. This was split into two 75k data sets, for training and for validation by our previously published labelling network Deep-Channel (Celik et al., 2020). (A) Shows the training and validation accuracy; the accuracy on the training set itself is near perfect, but it fails to an accuracy of 0.5 (to predict open or closed) with the unseen validation dataset. (B) Shows similar to (A), but with losses. Complete failure with the unseen validation set. Ergo, model training fails as is illustrated in (C) where the raw signal (top) is shown above the synchronised ground truth black and prediction red. In the second part of this demonstration (D, E and F), we replace the training data with the GAN generated dataset of Phenotype C, which could be any length, but in this example is 5x larger than the original. Now Deep-Channel does much better. After about 5 epochs, performance against the validation dataset improves in terms of both accuracy (D) and loss (E) and the final product is well trained model (F). There is close agreement between the ground truth (black) and the prediction (red).

The new GAN synthesis method will have strengths and weaknesses compared to stochastic Markovian model-based processes. (1) Speed: Training a GAN takes a considerable time (hours), but simulation itself takes ms per record. The usual stochastic methods produce records at the ms per record timescale, but do not need training. They still need laborious hand customising to simulate a particular phenotype of channel. Our software only Markovian based simulations, written in Python, take about 8ms to generate one “record” whereas DeepGANnel, on our GPU workstations takes about 4ms (once trained). It should be noted that Stochastic models will take n times longer to simulate data records with n channels within, whereas the GAN method would need retraining to a multi-channel dataset; but still take the same 4ms per record once trained. The method we used to create the training set in our DeepChannel project (Celik et al., 2020) used stochastic simulation followed by passing these data through a real patch clamp amplifier. This method was the slowest of all, producing data at less than real time (so approximately 1000x slower than other methods). (2) Authenticity: Speed is not the objective of this work however, it is “authenticity”. In the present paper, we provide metrics for authenticity in terms of UMAP/t-SNE cluster similarity for DeepGANnel, however there is no absolute way to do the same with stochastic methods. The more effort the user puts into it, analysing noise and reproducing these, measuring single channel properties and encoding this, the closer it would become, but this entire laborious process would need to be repeated for every type of ion channel and condition to be simulated. With DeepGANnel, the user simply points the script at a new set of seed data. Fundamentally, stochastic data may include unrecognised biases since every feature must be hand crafted, and DeepGANnel would include artefacts found in native data that would likely be omitted using a stochastic approach. The inclusion of these in training data would be important for development of *robust* analysis software. (3) A priori assumptions: DeepGANnel needs none, but stochastic simulation requires every detail to be estimated. However, stochastically generated data has the advantage that the experimenter could potentially develop software similar to HJCFit (Colquhoun, Hatton & Hawkes, 2003; Gibb et al., 2018) and QuB (Qin, Auerbach & Sachs, 1996), to recover the HMM, and validate accuracy. With a DeepGANnel simulated dataset the underlying HMM would be unknown, and need to be estimated with further software such as HJCFit, in the same way as one would do with real patch clamp data.

The literature includes previous examples where timeseries data can be simulated with a GAN (For example, Zhu, Ye, Fu, Liu & Shen, 2019), but these lack the synchronous labelling critical in single molecule analysis or many other physiological studies. We too found, in preliminary work (data not shown), that generating single-molecule timeseries with a basic GAN (Vanilla-GAN) model was very effective at producing authentic ion molecule signal patch-clamp signal, but this also lacked the output of the critical timepoint-by timepoint labelling necessary to meet our goals of creating valid alternative datasets. However, by switching to a 2DCNN based model in a shape of (samples x 2 × 1280 × 1) was very effective, with a small amount of carefully annotated seed data generating unlimited synthetic copies. In Figures 3 and 4 we show a direct comparison for a typical electrophysiological work up of both the original and DeepGANnel synthesised data. On the supplementary information and public repository, we include a movie of the training process. The match between synthetic kinetic analysis and the real data is not perfect, but rather close. The notable exception is that the longer states (both open and closed) are missing or have a diminished representation. We attribute this to the necessity to use a finite window (“image” width or “record length”) size. Also as stated in the methods this was cropped to remove leading and trailing artefacts. Perhaps models using far greater window lengths would be possible, but this does not appear to be a major problem for our purpose (since most single molecule events durations are within this window) and it would increase the model complexity many-fold. This model took 24-48 hours to train on our system and note that performance peaked but would deteriorate if it was left indefinitely. Our code allowed manual adjustment of learning rate as epochs progressed, but we still chose to stop the modelling manually. The ever-increasing GPU power make ever larger window sizes less of an issue in the future.

The potential for further exploitation, of GAN technology in electrophysiology beyond the current use for creation of datasets is immense. One future goal will be to simulate far more complex data signals, but we still have the limitation on how to acquire the fully annotated seed data in the first place. Potentially painstaking manual annotation of very short sections of data with known numbers of ion channel molecules by several human experts would be possible. Furthermore, it is possible that single-molecule GAN could be used more directly in electrophysiological modelling. Currently, single molecule behaviour within such models is derived by a set of differential equations based on a set of measured or even estimated parameters (Feetham, Nunn, Lewis, Dart & Barrett-Jolley, 2015), but it may be possible in the future to use GAN to generate more realistic stochastic behaviour directly. Additionally, future studies will investigate whether interpretability methods can be used to identify important, defining features within each different dataset that are missed either by eye or by standard single molecule analysis techniques.

The architecture we present here, using deep learning to generate physiological timeseries data with continuous annotations, could also be adapted easily for additional usability for equivalent systems in physiology. For example, action potential or electrocardiogram simulation. As proof-of-principle we show here that indeed DeepGANnel can easily synthesise telemetered ECG signal, again fully annotated (Figure 8). In this example the annotation dimension is merely beat (binary state 1) or no beat binary (0), but this could easily be extended to include P-wave (categorical state 2), T-wave (categorical state 3), or abnormal event (categorical state 4) etc with trivial code adaptation. Another example would potentially be Nanopore data which has similar data output to patch-clamp data.

**Fig. 8.**
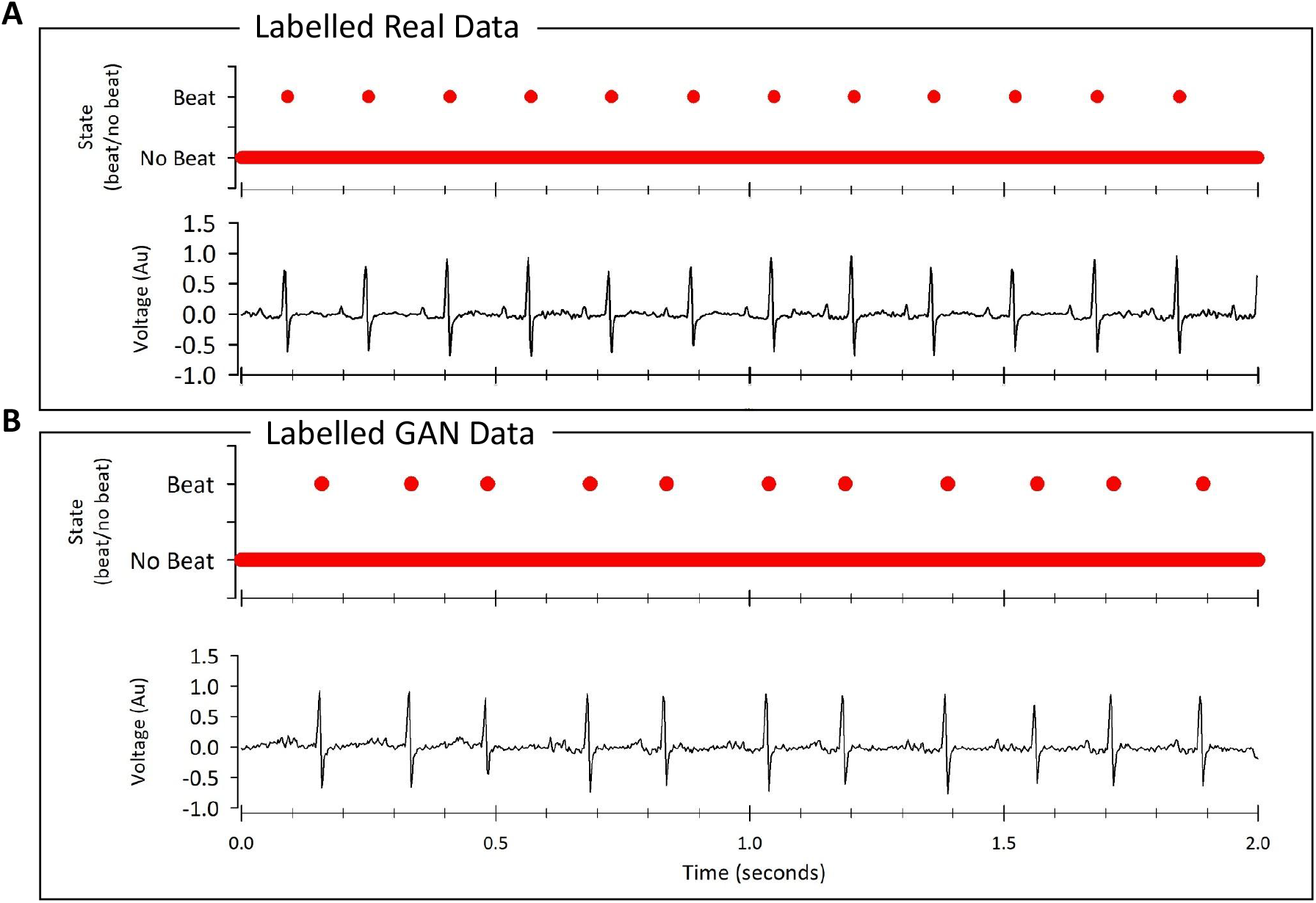
General usability of DeepGANnel for 2-dimensional physiological time series data. In this example we fed rodent ECG data (dimension 1) along with a beat annotation (dimension 2, peak of r-wave) into DeepGANel and trained with 10,000 epochs until realistic rodent ECG data was generated. Data collected via electrocardiogram transmitters from male Wistar rats (ETA-F20; Data Sciences International, St Paul, MN, USA) as previously described (Nunn, Feetham, Martin, Barrett-Jolley & Plagge, 2013), briefly ECG signal was digitized to a PC with a CED Micro1401 using Spike2 at 5 kHz. In principle this could also be encoded with further annotations such as t-wave (note rats do not show a significant p-wave).

In summary, GANs are increasingly proving a viable method to generate synthetic datasets for biological research, and here we show an implementation that allows simulation of time dependent single molecule (patch clamp ion channel protein) activity along with a continuous state annotation that is extendable for an array of physiological uses.

## 5. Acknowledgements

This work was funded by the BBSRC with grants: BB/R022143/1, BB/S008136/1 and a BBSRC DTP Studentship

## Appendix 1

Our neural networks are constructed by using so called “layers” of types of neruons. In this appendix we will go into substantially more detail as to how these “layers” work, and how the networks operate together during training. The full model architecture of both generator and discriminator including shapes are included in Appendix 2 and 3 respectively.

### Dense Layer

Dense layers, also known as “fully connected” layers, have each neuron connected to every neuron or data point of the previous layer (Figure 9). Let *x =* (*x*_1_,*x*_2_, *… x*_n_) be the inputs to the layer (either from the latent noise vector for the generator or the penultimate layer output for the discriminator). Then the output vector y of size m is given by:

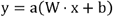

**Fig. 9.**
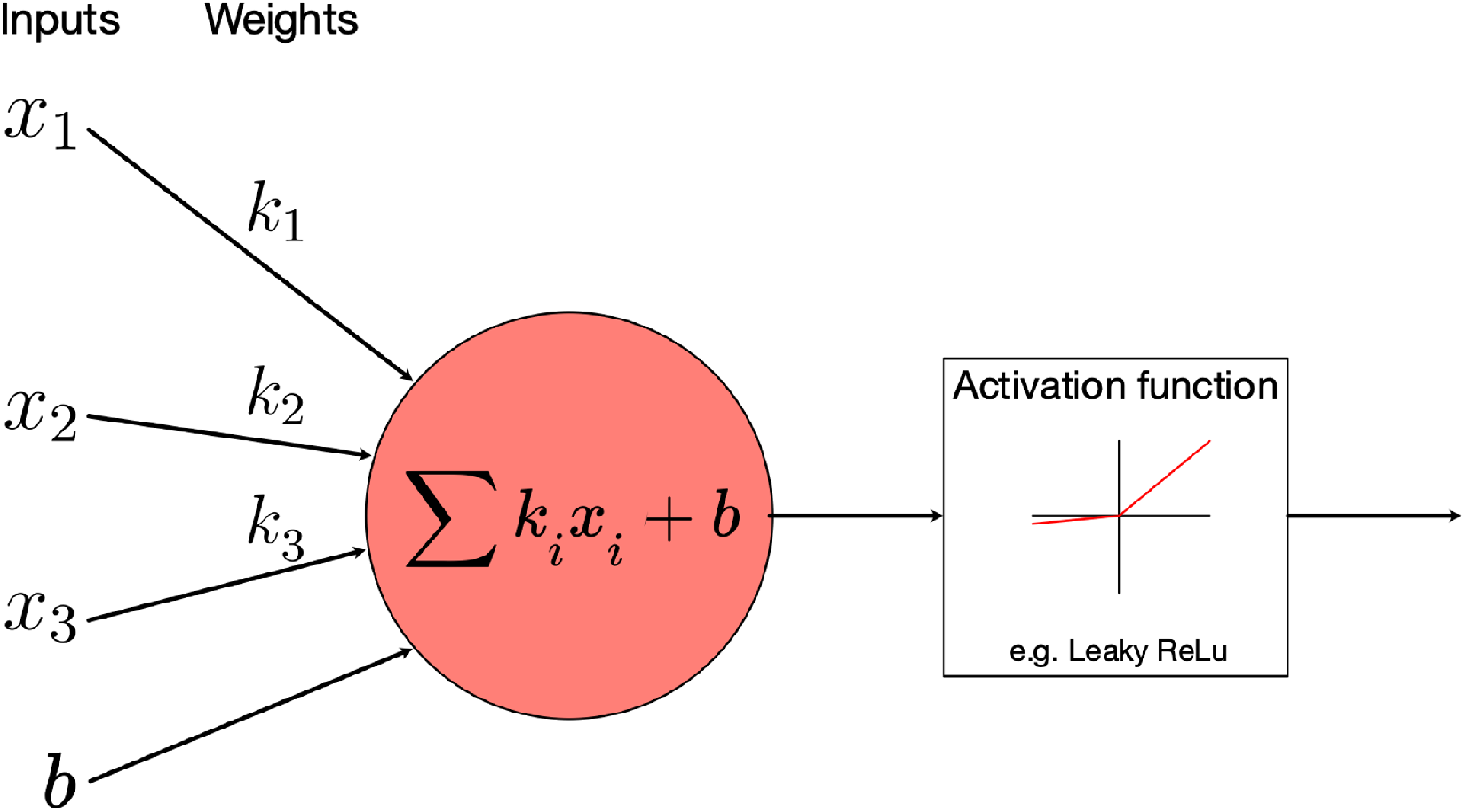
Diagram of how a simple neuron operates in a neural network. A number of inputs (either from an input vector or a series of outputs from other neurons) are multiplied by a series of trainable parameters. These are then totalled along with a non-trainable bias term and passed through to an activation function for output. The main disadvantage of using layers of simple neurons that have inputs from every previous output is parameter cost. As more neurons are added the number of parameters exponentially increases leading to poor training performance and the danger of overfitting.

Where W is the (n x m) matrix of weights that are adjusted throughout the training process by our optimizer funciton, b is the bias term, and a the activation function, which for all layers in both of our models is the Leaky Rectified Linear Unit (Leaky ReLU), that operates elementwise on the vector input by the following mechnaism:

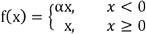

For all our functions, *α* is taken to be 0.3.

### Convolutional and Deconvolutional Layers

Convolutional and deconvolutional layers work via very similar means to achieve down-sampling and up-sampling respectively. Typically two dimensional layers are used for image analysis, but the principles can be extended to any number of dimensions.

At its core, the process involves using a kernel K of a chosen size with trainable parameters that passes over the input and is multiplied by a sub-matrix of the input X - the size of this submatrix, and the order of operations dictates whether downsampling or upsampling occurs.

In downsampling, each element y_i,j_ of the output matrix Y is given as follows - for simplicity here the kernel size is (3 × 3) and the stride is 1. Figure 10 shows visualisations of the areas chosen and the mappings. they correspond to.

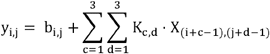

**Fig. 10.**
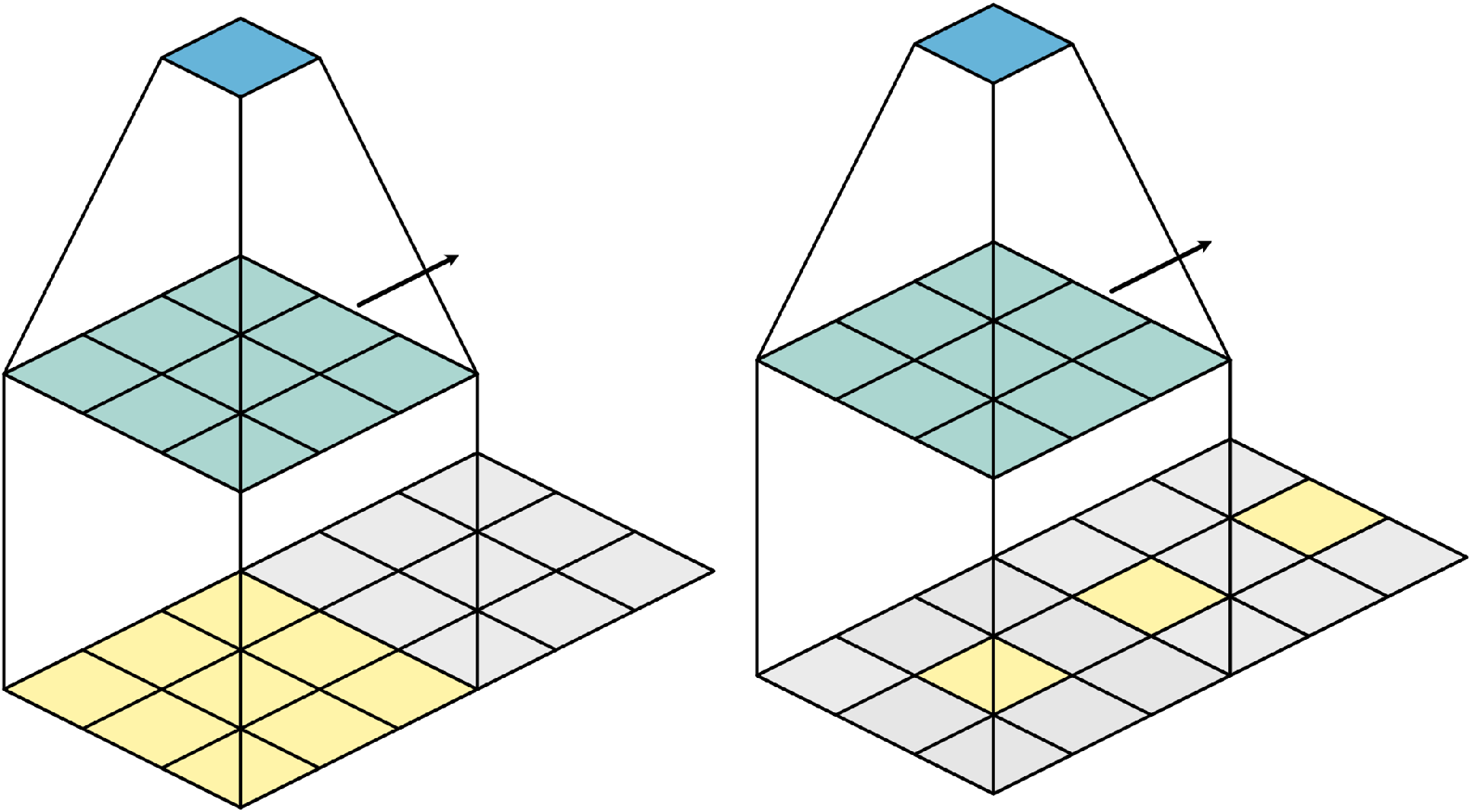
Diagram of Convolutional and deconvolutional layers. Typically used in image analysis, convolutional (left) and deconvolutional layers (right) work by passing a series of trainable kernels (green) over an image, taking the sum of the pairwise product of the kernel and each submatrix of the input. As these kernels “scan” across the image, features such as edges are detected, and “feature maps” are built as the network increases in depth. By adding zero padding between each input, the downscaling effect of the scanning process becomes upscaling, which can be used in generation instead of detection.

The dimensions of the kernel are adjusted by changing the bounds of the sums, and the indexes of the input matrix (the -1 term becomes -⌊s/2⌋, where s is the desired kernel size). A stride can be applied in either or both directions by incrementing i and/or j in values other than one, and simply ignoring the resulting “gaps” in the output matrix. Typically when either of the indexes of X are outside of the bounds of X (for example, when the index is negative), the value is taken as zero (this is refered to as padding).

### Max Pooling Layers

Max pooling layers achieve down sampling but are far simpler than convolutional layers. A set of n x n submatricies are taken, and for each one the maximal value of each submatrix is passed to a new, smaller submatrix. For example, when n = 2, the output matrix will be of size 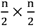. This quickly and cheaply achieves downsampling but does not recognise patterns as strongly as convolutional layers.

### Dropout Layers

Droupout layers are a regularisation technique designed to stop a model overfitting. They work by disabling a randomly selected proportion of the inputs, with the proportion being defined with a probability hyperparameter p.

### Batch Normalisation

Batch normalisation is another regularisation technique. In theory, a layer’s parameters are adjusted based on the assumption that the previous layer’s parameters are static; however this is clearly not the case. A batch normalisation layer scales the output of a layer to have a mean of 0 and standard deviation of 1 for each batch to realign the data with this assumption.

## Notes

https://github.com/RichardBJ/DeepGannel

### Competing Interest Statement

The authors have declared no competing interest.

### Summary of Updates

Major revision is inluding extra channels and manifold/clustering comparions

## References

Anderson DF, Ermentrout B, & Thomas PJ (2015). Stochastic representations of ion channel kinetics and exact stochastic simulation of neuronal dynamics. Journal of Computational Neuroscience 38: 67–82.

Brock A, Donahue J, & Simonyan K (2019). Large Scale GAN Training for High Fidelity Natural Image Synthesis. 1809.11096.

Celik N, O’Brien F, Brennan S, Rainbow RD, Dart C, Zheng Y, et al. (2020). Deep-Channel uses deep neural networks to detect single-molecule events from patch-clamp data. Commun Biol 3: 3.

Colquhoun D, Hatton CJ, & Hawkes AG (2003). The quality of maximum likelihood estimates of ion channel rate constants. J Physiol 547: 699–728.

Colquhoun D, Hawkes AG, & Srodzinski K (1996). Joint distributions of apparent open and shut times of single-ion channels and maximum likelihood fitting of mechanisms. Philosophical Transactions of the Royal Society of London Series A: Mathematical, Physical and Engineering Sciences 354: 2555–2590.

Delaney AM, Brophy E, & Ward TE (2019). Synthesis of Realistic ECG using Generative Adversarial Networks. 1909.09150.

Donahue C, McAuley J, & Puckette M (2019). Adversarial Audio Synthesis. 1802.04208v3.

Feetham CH, Nunn N, Lewis R, Dart C, & Barrett-Jolley R (2015). TRPV4 and KCa ion channels functionally couple as osmosensors in the paraventricular nucleus. Br J Pharmacol 172: 1753–1768.

Gibb AJ, Ogden KK, McDaniel MJ, Vance KM, Kell SA, Butch C, et al. (2018). A structurally derived model of subunit-dependent NMDA receptor function. J Physiol 596: 4057–4089.

Gillespie DT (1977). Exact Stochastic Simulation of Coupled Chemical-Reactions. J Phys Chem-Us 81: 2340–2361.

Gnanasambandam R, Nielsen MS, Nicolai C, Sachs F, Hofgaard JP, & Dreyer JK (2017a). Unsupervised Idealization of Ion Channel Recordings by Minimum Description Length: Application to Human PIEZO1-Channels. Frontiers in Neuroinformatics 0: 31–31.

Gnanasambandam R, Nielsen MS, Nicolai C, Sachs F, Hofgaard JP, & Dreyer JK (2017b). Unsupervised idealization of ion channel recordings by minimum description length: application to human PIEZO1-channels. Frontiers in neuroinformatics 11: 31.

Goodfellow IJ, Pouget-Abadie J, Mirza M, Xu B, Warde-Farley D, Ozair S, et al. (2014). Generative Adversarial Nets. Advances in Neural Information Processing Systems 27 (NIPS 2014).

Gretton A, Borgwardt K, Rasch M, Schölkopf B, & Smola AJ (2008). A kernel method for the two-sample-problem. 0805.2368.

Gretton A, Borgwardt KM, Rasch MJ, Scholkopf B, & Smola A (2012). A Kernel Two-Sample Test. J Mach Learn Res 13: 723–773.

Hamill OP, Marty A, Neher E, Sakmann B, & Sigworth FJ (1981). Improved Patch-Clamp Techniques for High-Resolution Current Recording from Cells and Cell-Free Membrane Patches. Pflugers Archiv-European Journal of Physiology 391: 85–100.

Hodgkin AL, & Huxley AF (1952). The components of membrane conductance in the giant axon of Loligo. J Physiol: 473-496.

Hotz T, Schutte OM, Sieling H, Polupanow T, Diederichsen U, Steinem C, et al. (2013). Idealizing Ion Channel Recordings by a Jump Segmentation Multiresolution Filter. IEEE Transactions on NanoBioscience 12: 376–386.

Imbrici P, Conte Camerino D, & Tricarico D (2013). Major channels involved in neuropsychiatric disorders and therapeutic perspectives. Frontiers in Genetics 4: 76.

Juette MF, Terry DS, Wasserman MR, Altman RB, Zhou Z, Zhao H, et al. (2016). Single-molecule imaging of non-equilibrium molecular ensembles on the millisecond timescale. Nature methods 13: 341.

Karras T, Laine S, & Aila T (2019). A Style-Based Generator Architecture for Generative Adversarial Networks. 1812.04948v3.

Lewis R, Feetham CH, Gentles L, Penny J, Tregilgas L, Tohami W, et al. (2013). Benzamil sensitive ion channels contribute to volume regulation in canine chondrocytes. Brit J Pharmacol 168: 1584–1596.

McInnes L, Healy J, & Melville J (2018). UMAP: Uniform Manifold Approximation and Projection for Dimension Reduction. 1802.03426.

Mobasheri A, Lewis R, Maxwell JEJ, Hill C, Womack M, & Barrett-Jolley R (2010). Characterization of a stretch-activated potassium channel in chondrocytes. Journal of Cellular Physiology 223: 511–518.

Neher E, & Sakmann B (1976). Single-Channel Currents Recorded from Membrane of Denervated Frog Muscle-Fibers. Nature 260: 799–802.

Nicolai C, & Sachs F (2013). Solving ion channel kinetics with the QuB software. Biophysical Reviews and Letters 8: 191–211.

Nunn N, Feetham CH, Martin J, Barrett-Jolley R, & Plagge A (2013). Elevated blood pressure, heart rate and body temperature in mice lacking the XLαs protein of the Gnas locus is due to increased sympathetic tone. Exp Physiol 98: 1432–1445.

Qin F (2004). Restoration of Single-Channel Currents Using the Segmental k-Means Method Based on Hidden Markov Modeling. Biophysical Journal 86: 1488–1488.

Qin F, Auerbach A, & Sachs F (1996). Estimating single-channel kinetic parameters from idealized patch-clamp data containing missed events. Biophysical Journal 70: 264–280.

Qin X-H, Wang Z-Y, Yao J-P, Zhou Q, Zhao P-F, Wang Z-Y, et al. (2020). Using a one-dimensional convolutional neural network with a conditional generative adversarial network to classify plant electrical signals. Computers and Electronics in Agriculture 174: 105464.

Quinton PM (1990). Cystic fibrosis: a disease in electrolyte transport; Cystic fibrosis: a disease in electrolyte transport. The FASEB Journal 4: 2709–2710.

Radford A, Metz L, & Chintala S (2015). Unsupervised representation learning with deep convolutional generative adversarial networks. arXiv preprint 151106434.

Sakmann B, & Neher E (1984). Patch Clamp Techniques for Studying Ionic Channels in Excitable Membranes. Annu Rev Physiol 46: 455–472.

Sakoe H, & Chiba S (1978). Dynamic-Programming Algorithm Optimization for Spoken Word Recognition. Ieee T Acoust Speech 26: 43–49.

Salvadora S, & Chan P (2007). Toward accurate dynamic time warping in linear time and space. Intell Data Anal 11: 561–580.

Serviss JT, Gådin JR, Eriksson P, Folkersen L, & Grandér D (2017). ClusterSignificance: a bioconductor package facilitating statistical analysis of class cluster separations in dimensionality reduced data. Bioinformatics 33: 3126–3128.

Silver KS, D. Y, Nomura Y, Oliveira EE, Salgado VL, Zhorov BS, et al. (2014). Voltage-Gated Sodium Channels as Insecticide TargetsElsevier, pp 389–433.

Skerratt SE, & West CW (2015). Ion channel therapeutics for pain. Channels 9: 344–351.

Truong ND, Kuhlmann L, Bonyadi MR, Querlioz D, Zhou L, & Kavehei O (2019). Epileptic seizure forecasting with generative adversarial networks. IEEE Access 7: 143999–144009.

Van Der Maaten L, & Hinton G (2008). Visualizing Data using t-SNE. J Mach Learn Res 9: 2579–2605.

Voldsgaard Clausen M (2020). Obtaining transition rates from single-channel data without initial parameter seeding. Channels (Austin) 14: 87–97.

Welsh MJ (1990). Abnormal regulation of ion channels in cystic fibrosis epithelia; Abnormal regulation of ion channels in cystic fibrosis epithelia. The FASEB Journal 4: 2718–2725.

Zhu F, Ye F, Fu Y, Liu Q, & Shen B (2019). Electrocardiogram generation with a bidirectional LSTM-CNN generative adversarial network. Sci Rep 9: 6734.

