## Supplementary figures and images for "DeepGANnel: Synthesis of fully annotated single molecule patch-clamp data using generative adversarial networks"

### Appendix 2

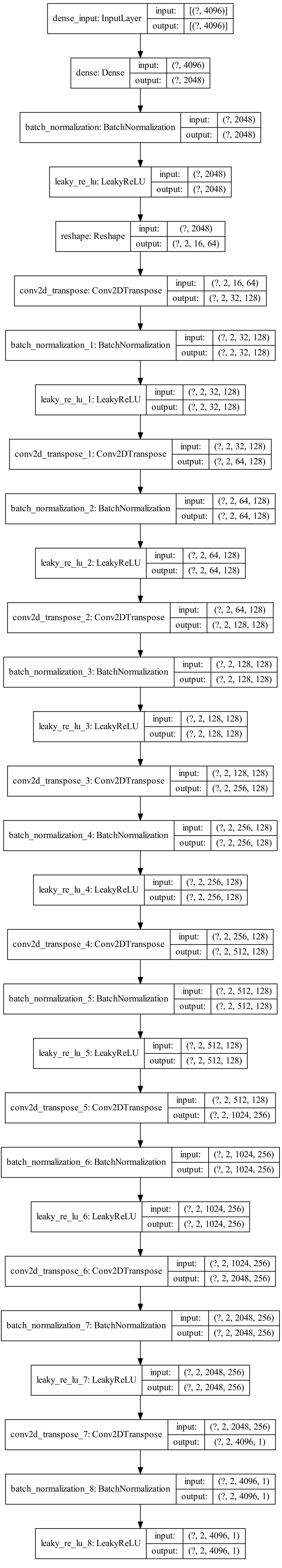

### Appendix 3

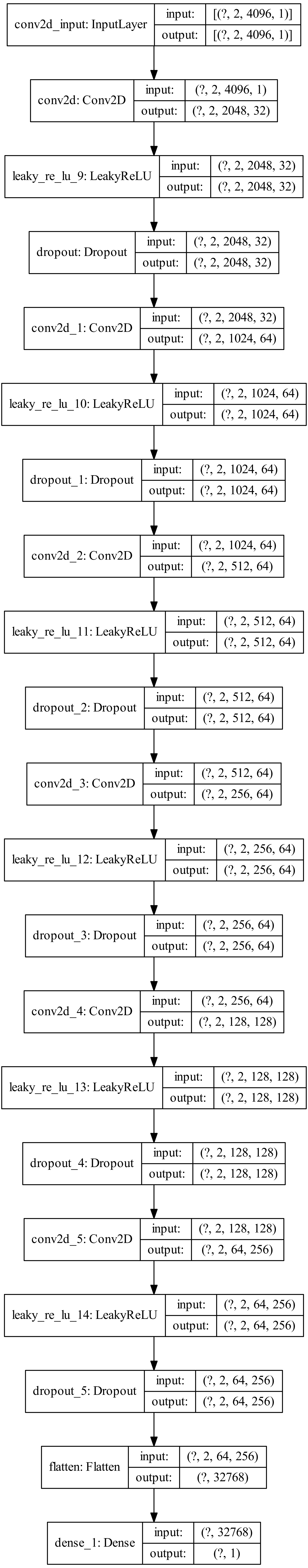
